# DeepInterface: Protein-protein interface validation using 3D Convolutional Neural Networks

**DOI:** 10.1101/617506

**Authors:** A.T. Balci, C. Gumeli, A. Hakouz, D. Yuret, O. Keskin, Attila Gursoy

## Abstract

**Motivation:** Protein–protein interactions are crucial in almost all biological processes. Proteins interact through their interfaces. It is important to determine how proteins interact through interfaces to understand protein binding mechanisms and to predict new protein-protein interactions.

**Results:** We present DeepInterface, a deep learning based method which predicts, for a given protein complex, if the interface between the proteins of a complex is a true interface or not. The model is a 3-dimensional convolutional neural networks model and the positive datasets are obtained from all complexes in the Protein Data Bank, the negative datasets are the incorrect solutions of the docking decoys. The model analyzes a given interface structure and outputs the probability of the given structure being an interface. The accuracy of the model for several interface data sets, including PIFACE, PPI4DOCK, DOCKGROUND is approximately 88% in the validation dataset and 75% in the test dataset. The method can be used to improve the accuracy of template based PPI predictions.

## 1 Introduction

Protein–protein interactions are crucial in almost all biological processes. Proteins interact through their interfaces on their surfaces (Keskin, et al., 2008). Interfaces are made of interacting residues that come from two interacting proteins. In order to know the interfaces, we have to have the three dimensional structures of the protein-protein complexes. However, only a small fraction of all interactions have this information. Computationally, docking techniques are frequently used for protein-protein structure prediction (Lensink, et al., 2016). Once docking is run for two proteins, hundreds or thousands of solutions called decoys are obtained. Therefore, it is important to be able to discriminate the native-like solutions from the incorrect predictions of docking experiments, one way to succeed this is to be able to predict accurately the interface regions of interacting proteins.

Interfaces may consist of some continuous segments and some single residues from each of the proteins (Tuncbag, et al., 2008; Winter, et al., 2012). Interface properties that can distinguish interface residues from the rest of the protein surface residues include the shape of the binding interfaces, ranges of surface areas, amino acid types and polar/nonpolar nature of interface residues, the hydrogen bonds and salt bridges across the interfaces, residue conservation, hot spots and the types of secondary structures in the interface region, phosphorylation and methylation status of residues (Jones and Thornton, 1997; Keskin, et al., 2008; Keskin, et al., 2008; Li, et al., 2004; Tuncbag, et al., 2008).

These properties are used a priori towards prediction (Aytuna, et al., 2005; Baspinar, et al., 2014; Ogmen, et al., 2005; Tuncbag, et al., 2011), allowing us to focus on the binding sites, or can be used a posteriori for scoring/ranking the predicted complexes (Liang, et al., 2009; Nadalin and Carbone, 2018) (Andreani, et al., 2013; Keskin, et al., 1998; Popov and Grudinin, 2015).

There are recent studies using interface structures to score protein-protein complexes. Nadalin and Carbone proposed CIPS (Combined Interface Propensity for decoy Scoring), a new pair potential combining interface composition with residue–residue contact preference (Nadalin and Carbone, 2018). Popov and Grudinin (Popov and Grudinin, 2015) demonstrated that knowledge of only native protein-protein interfaces is sufficient to construct well-discriminative predictive models for the selection of binding candidates using statistical learning. (Chen and Zhou, 2005) is a consensus neural network method for predicting protein-protein interaction sites. Given the structure of a protein, cons-PPISP will predict the residues that will likely form the binding site for another protein.

Growth of protein data bank and the accumulation of computational docking results provide a large amount of protein-protein interface structural data. These large data can be used in machine learning. A number of datasets of protein-protein interfaces have been extracted so far. These datasets list all available interfaces like ProtinDB (Jordan, et al., 2012), ProtCID (Xu and Dunbrack, 2011) or group interfaces into similar clusters, like PIFACE (Cukuroglu, et al., 2014). Unique interface structures can help as templates to model protein complexes, and according to that the existing interfaces already can achieve this aim (Baspinar, et al., 2014; Kundrotas, et al., 2012; Tuncbag, et al., 2011). With these large datasets, machine learning, especially deep learning in recent years has been applied to protein-protein interaction prediction, protein fold classification, and protein residue contact/folding using sequence and structure info. In general, machine learning based methods use higher-level features computed from sequence and structure data. One of the advantages of deep learning is the automatic feature extraction from raw data.

Our method, DeepInterface, is the first deep learning approach (to the best of our knowledge) utilizing a) only raw protein interface structure at atomic level and b) large set of interface data (around 270000 structures) for classifying a given predicted protein-protein interface as native or not.

Convolutional Neural Networks (CNN), a type of deep learning, have been very successful in image processing area and recently are being used for protein structure predictions. Most CNN applications for protein structure and fold determination used 1D or 2D CNNs with reduced input representations from protein sequence and structure. For example, DNCON2 is a protein residue-residue contact predictor (Adhikari, et al., 2018). As input it uses predicted solvent accessibility, secondary structure values, and coevolution features derived from multiple sequence alignment.. DeepContact (Liu, et al., 2018) is another protein residue-residue contact predictor where 1D and 2D protein structures are derived from protein sequence and structure. Protein structure and fold determination are not the only problem CNNs are applied. DeepFold (Liu, et al., 2018), for example is a protein structure motif extractor to be used for searching similar protein structures. It uses 2D distance matrices as the raw input derived from protein 3D structures. The structural motifs learned by the neural networks are used represent a fingerprint to represent protein structure.

The CNN studies mentioned above do not use 3D CNN and do not use raw protein structure, but use computed features from protein sequences. The most relevant approaches to ours in the way that used 3D protein structures as raw input and used 3D CNN are DeepSite (Jimenez, et al., 2017) and 3DCNN (Torng and Altman, 2017). DeepSite focuses on small molecule-protein interaction. It predicts binding sites using 3D CNNs. The protein structure is represented as a 3D grid of 1×1×1 A voxels. Voxels are labeled with pharmacophoric properties of the occupying atoms based on the coordinates and each label is used as a channel in 3D CNN network. 3DCNN uses 3D CNN and protein structure to learn the environment of individual amino acids to be used in identifying functionally important amino acids. The protein microenvironments (local regions) are represented as a 3D grid of size 20A around a central atom with four atom channels.

Our approach, DeepInterface, also uses 3D protein structure as raw data represented as a 25×25×25 3D grid covering the protein-protein interface and 40 channels based on amino acid types of two interacting proteins in the interface area. None of the above approaches focuses on protein-protein interfaces. DeepInterface differs also in the way raw data are represented: 20 channels based on coordinates of atoms and residue type rather than pharmacophoric properties or atom types are used. DeepInterface is trained with considerable amount of raw data, around 270000 protein-protein interface structure gathered from Protein Data Bank (PDB) and from docking decoy sets DOCKGROUND (Liu, et al., 2008)and PPI4DOCK (Yu and Guerois, 2016).

## 2 Materials and Methods

### 2.1 Data

We collected data for training and testing our model from 4 sources: DOCKGROUND (Liu, et al., 2008) PPI4DOCK(Yu and Guerois, 2016), PIFACE(Cukuroglu, et al., 2014) and PDB. The most of the positive interfaces are taken from PIFACE which is a collection of interface structures extracted from PDB files published until 2012. We obtained the rest of our positive dataset from the PDB deposited after 2012 to avoid duplicates. The other two data sets, DOCKGROUND and PPI4DOCK decoy sets, provide the negative interfaces.

Before explaining the data preparation and data sets, what is meant by “positive” and “negative” interface should be clarified. An interface in a protein complex is the set of contacting or nearby residues. Two residues belonging to different chains are in contact if any pair of heavy atoms from the two residues are within their vdW distance plus 0.5 A (as defined in PRISM (Baspinar, et al., 2014)). A residue is a nearby residue if the distance between CAs of it and a contacting residue is less than 6 A. We consider an interface to be positive if it comes from a complex deposited in PDB. If an interface comes from a docking complex, to determine whether it is is negative or not, we used the classification developed by Critical Assessment of Predicted Interactions (CAPRI) (Janin, et al., 2003). Within the scope of CAPRI, predicted complexes are evaluated into 4 categories based on 3 quantities: Fnat (fraction of native contacts), LRMSD (ligand root mean square deviation), and iRMSD (interface root mean square deviation). If Fnat of a complex is lower than 0.1 or its iRMSD is greater than 10A while its iRMSD is greater than 4A, the complex is classified as “incorrect”. We consider interfaces obtained from “incorrect” complexes as negative interfaces.

We selected around 130000 negative interfaces from the PPI4DOCK benchmark. PPI4DOCK has 1417 unbound models which can produce up to 54000 structures each. PPI4DOCK provides CAPRI assessment of each structure. These models, consisting of two chains, are heterodimers. To create structures from the models, ZDOCK (Chen, et al., 2003) docking tool is used. ZDOCK being an FFT based algorithm exhaustively tries all possible translations and rotations of the ligand unit on the receptor unit with a unit step. Taking advantage of the exhaustive search carried out by ZDOCK, we selected approximately 110 incorrect structures with the ligand being in different positions from each model. We tried to avoid similar or highly overlapping interfaces while choosing 110 incorrect structures. After the selection, the interfaces are extracted as described above.

DOCKGROUND is an unbound benchmark very similar to PPI4DOCK. It has two separate decoy sets. Set 1 consists of 61 complexes while Set 2 consists of 351 complexes. To create structures, another FFT based tool GRAMM-X (Tovchigrechko and Vakser, 2006) is used. Each complex in both sets have 100 structures at least one of which is near native (lRMSD < 5A). DOCKGROUND benchmark provides lists of LRMSD, IRMSD, and Fnat values for each structure. We selected structures satisfying CAPRI’s incorrect criteria and applied the same interface extraction procedure above.

The PIFACE dataset is our main source of positive interfaces. The dataset consists of approximately 130000 interfaces extracted from approximately 80000 PDB files. We included already extracted interfaces from PIFACE in our positive dataset. To enrich our positive dataset, we also extracted 8125 interfaces from the PDB files which are published after 2012.

After the preparation of the interfaces from all the data sets, we ended up with 271,830 interfaces to train and test our model. 135,915 interfaces in the database are positive while the other half is negative. We separated PIFACE and PPI4DOCK to train our models, and DOCKGROUND and PDB files published after 2012 to test our models. The reason behind the separation is to see how well our model generalizes to structures coming from different sets. To summarize, our training and validation data set comprises 127790 negative interfaces coming from PPI4DOCK and 127790 positive interfaces coming from PIFACE, our test data set comprise 8125 positive interfaces from PDB files published after 2012, and 8125 negative interfaces coming from DOCKGROUND.

After the interface extraction step, we rotated all interfaces to give them similar poses, since CNNs are not rotation invariant, We translated the midpoints between the centers of masses of the two sides of the interfaces to the origin and rotated the centers of mass onto x axis. We also rotated the interfaces so that the atom which is furthest from the origin, after the first transformation, points to positive z direction. After the rotation and translation steps, 40 by 25 by 25 by 25 matrices are created for every interface. The CNN model is designed to work on 3 dimensional matrices with 40 channels: one channel each for the side chains of the each chains’ every 19 residue types excluding GLY and 1 channel each for the alpha Cs (**Fig 1** left panel). The shape is determined so that every cell represents 2×2×2 A^3^ of 50×50×50 A^3^ of space which includes the interface. If an interface cannot fit into 50 ×50×50 A^3^ with its given position, that interface is eliminated from the data set. The amino acids are considered as 3D Gaussian distributions with the coordinates of C-alpha and C-beta as the means. The standard deviation of the distributions are derived from the radii of the each amino acid types. The distributions are quantized to fill cells of matrices (as shown in **Fig 1** right panel).

**Fig 1.**
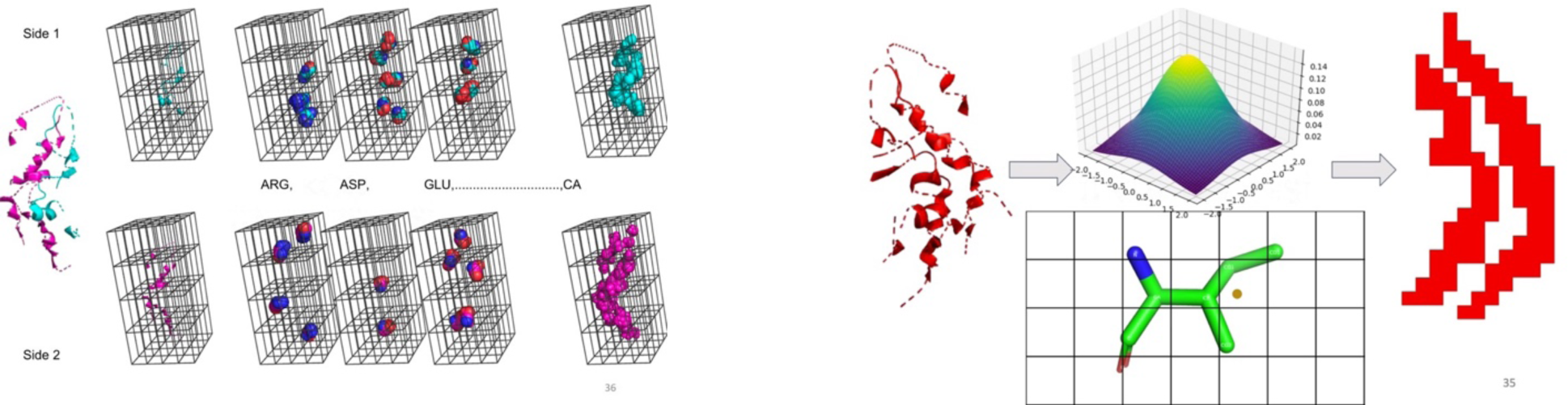
(Left panel) Data Preparation for CNN 40 channels. For one side of the interface, 19 channels for each amino acid type except for GLY, and 1 channel for Cα. (Right panel) - Grid representation of atoms.

### 2.2 DeepInterface Convolutional Neural Network

Our network is composed of 4 convolutional layers followed by a global pooling layer and 2 fully connected layers. All convolutional layers have 128 5×5×5 filters, with stride and padding set to 2. Each convolution is followed by a batch normalization (Ioffe and Szegedy, 2015)and rectifier linear unit (ReLU) activation (Nair and Hinton, 2010). We employed a global average pooling at the end of convolutional layers for improving generalization, which converts the 128 4×4×4 spatial feature maps to 128 global features. Global pooling is followed by a fully connected layer with 512 hidden units, batch normalization and ReLU. The final layer outputs two score values for negative and positive classes (see **Fig 2**).

**Fig 2.**
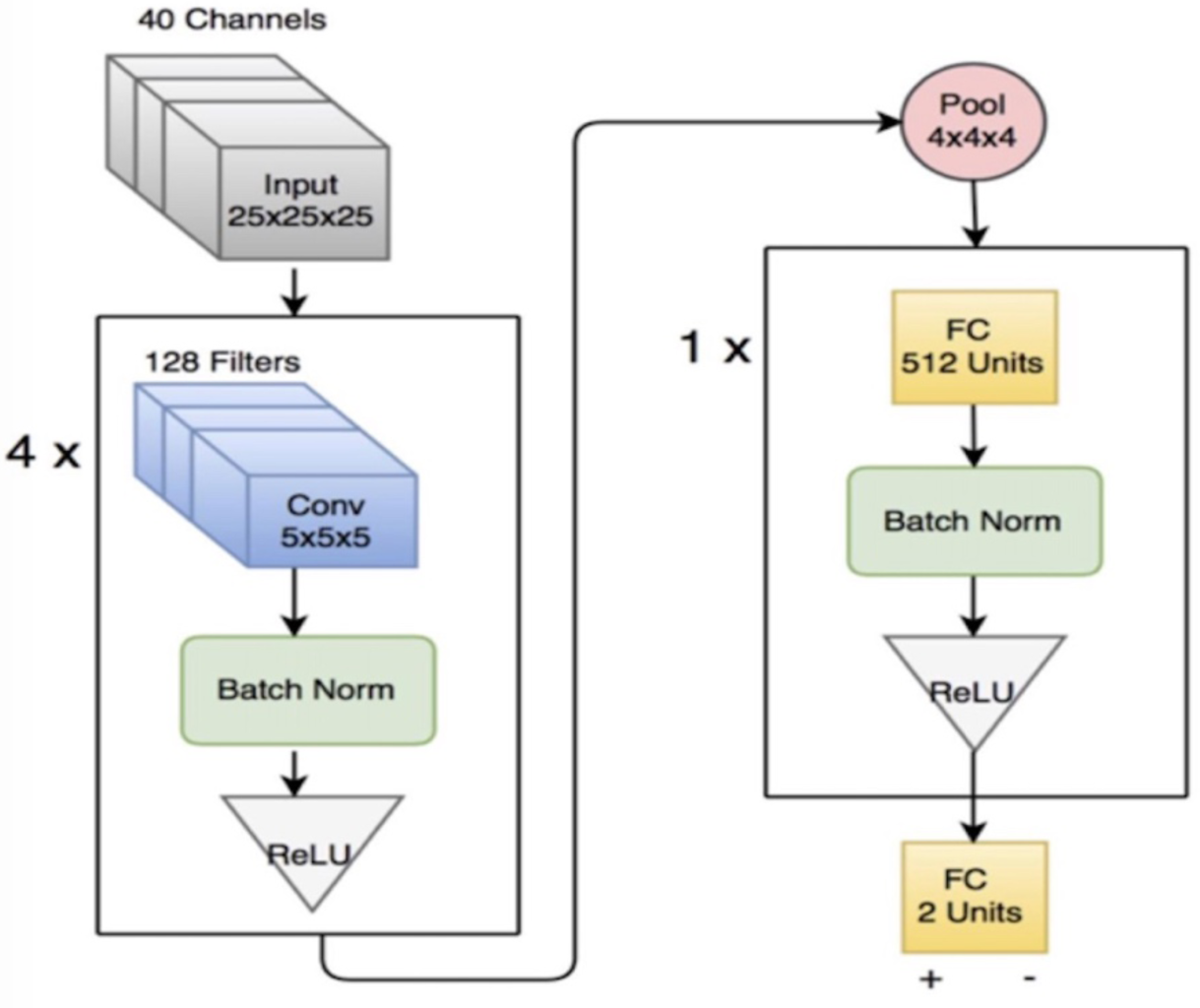
The schematics diagram of the DeepInterface CNN.

Aggressively decreasing spatial dimensions with strided convolutions while keeping the channel size constant is to address sparsity of our data. Global pooling considerably reduces the number of parameters for better generalization. Finally, having a fully connected hidden layer at the output allows our model to have a better decision-making capability using non-linearity.

In our experiments, we observed that this simple architecture has better convergence properties and is less prone to over-fitting compared to both classical Lenet-like models (Lecun, et al., 1998) and very deep residual architectures (He, et al., 2016)). The model is developed using the KNet (Yuret, 2016), a deep learning framework implemented in Julia (Bezanson, et al., 2017).

### 2.3 Training Method

In our experiments, we used Adam optimizer (Kingma and Ba, 2014) with learning rate 0.001 and minibatch size of 32. Learning rate is decayed by a factor of 2 whenever validation loss increases compared to previous epoch. Training data is shuffled at the beginning of each epoch. We did not perform any dropout and weight decay for regularization. We initialized weights randomly, using kaiming normal distribution (He, et al., 2015) for convolutional layers and xavier uniform distribution (Xavier and Yoshua, 2010) for fully connected layers.

## 3 Results

### Chemical composition of the datasets

We first compared the amino acid composition of our four datasets, namely, the negative datasets from DOCKGROUND and PPI4Dock; and positive data sets from PIFACE and PDB2012. We grouped the amino acids into 7 groups according to their chemical properties. These groups are; aliphatic group, hydroxyl group, sulfur group, acidic group, basic group, aromatic group, imino group. We classified GLY, ALA, VAL, LEU, and ILE into the aliphatic group; SER, THR, TYR into the hydroxyl group; CYS, MET into the sulfur group; ASP, ASN, GLU, and GLN into the acidic group; LYS, ARG, HIS into the basic group; PHE, TYR, TRP Into the aromatic group; and finally, PRO into the imino group. We derived the distribution of percentages of each group for interfaces in each dataset. Almost all of the distributions are similar for all datasets. This result highlights that there is no bias in our negative decoys with respect to chemical compositions. We have to note that positive interfaces come from almost 130,000 crystal or NMR complexes, whereas the negative complexes are generated by using around 1,500 monomer structures. The most discerning feature comes from the aromatic and aliphatic groups. A representative distribution is provided in **Fig 3** for aromatic compositions. This figure shows that interfaces of the complexes in PPI4DOCK dataset have slightly a higher composition in aromatic residues compared to the rest.

**Fig 3.**
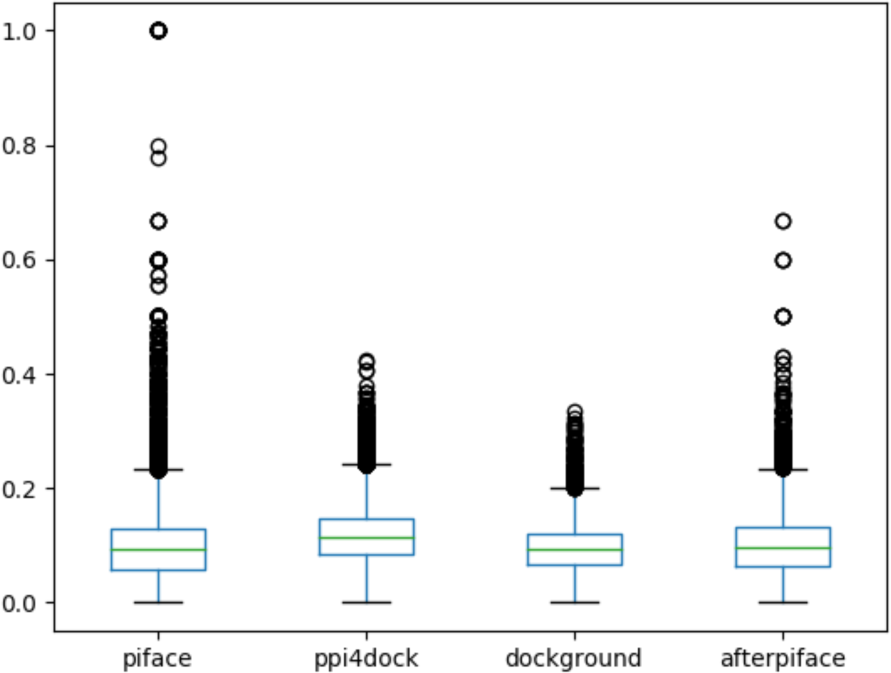
Distributions of aromatic compositions in four datasets.

### Performance and accuracy of DeepInterface

We trained the DeepInterface on PIFACE (positive) and PPI4DOCK (negative) datasets of 255580 interfaces. The datasets were uniformly sampled to form the validation set to size 35000 examples to evaluate the model during the training. After three epochs of training, the average validation accuracy converged to 88%. We experimented with larger splits and made sure the performance is consistent among different validation splits obtained in the same way.

Our CNN model provides two scores for each interface complex, one score for being a positive prediction, one score for being a negative prediction. The decision for a native or incorrect interface is made by computing the difference between the positive and negative scores (diff-score). **Fig 4** shows the distribution of the scores of the decoys of an interface (incorrect complexes) for two cases (2pukAB and 3c0aAB). Notice that, the negative scores are bigger than the positive ones for most decoys as expected. The black bar indicates the diff-score of the true interface. The yellow distribution highlights the frequency of native-like predictions (if the diff-score is larger than zero) or incorrect interface predictions (if the diff-score is smaller than zero). The model performed better for 3c0aAB in this example. We provide these cases to explain how we assign a single score to a prediction. The performance for all interfaces is given in Table 2 and Table 3. Table 2 provides the performance for the training, validation and testing data separately. The accuracy for the training is 0.92, for the validation is 0.88, and for the testing (unseen data) is 0.75. A recent review lists the comparison of various protein interface prediction methods (around 60 methods). The accuracies range between 85.4% and 45.0% - see Table 2 of (Zhao, et al., 2011). However, these methods used very different data sets and measurements, therefore it is difficult to compare one with the other directly. Our accuracy results for the validation (0.88) and the test (0.75) are well above the average of the reported results. In all cases, recall values are high meaning that we correctly identify the positives, however specificity values are lower indicating that we do not correctly predict the negatives.

**Table 1:** Dataset

|  | PIFACE | PPI4DOCK | DOCKGROUND | PDB2012 | Total |
| --- | --- | --- | --- | --- | --- |
| Training | 110,248 | 110,332 |  |  | 220,580 |
| Validation | 17,542 | 17,458 |  |  | 35,000 |
| Test |  |  | 8,125 | 8,125 | 16,250 |
| Total | 127,790 | 127,790 | 8,125 | 8,125 | 271,830 |

**Table 2:** Validation Accuracies of Our Best Models.

|  | Precision | Recall | Specificity | NPV | Accuracy |
| --- | --- | --- | --- | --- | --- |
| Training | 0.88 | 0.96 | 0.88 | 0.96 | 0.92 |
| Validation | 0.85 | 0.93 | 0.83 | 0.92 | 0.88 |
| Testing data | 0.61 | 0.87 | 0.68 | 0.90 | 0.75 |

**Table 3:** Ranking of the true interface among the decoys of the corresponding complex.

|  | All sets<br>(599 cases) | Training + Validation<br>(454 cases) | Validation + Testing<br>(32 cases) |
| --- | --- | --- | --- |
| Top1 | 0.41 | 0.49 | 0.22 |
| Top10 | 0.76 | 0.73 | 0.41 |
| Top20 | 0.86 | 0.86 | 0.59 |

**Figure 4.**
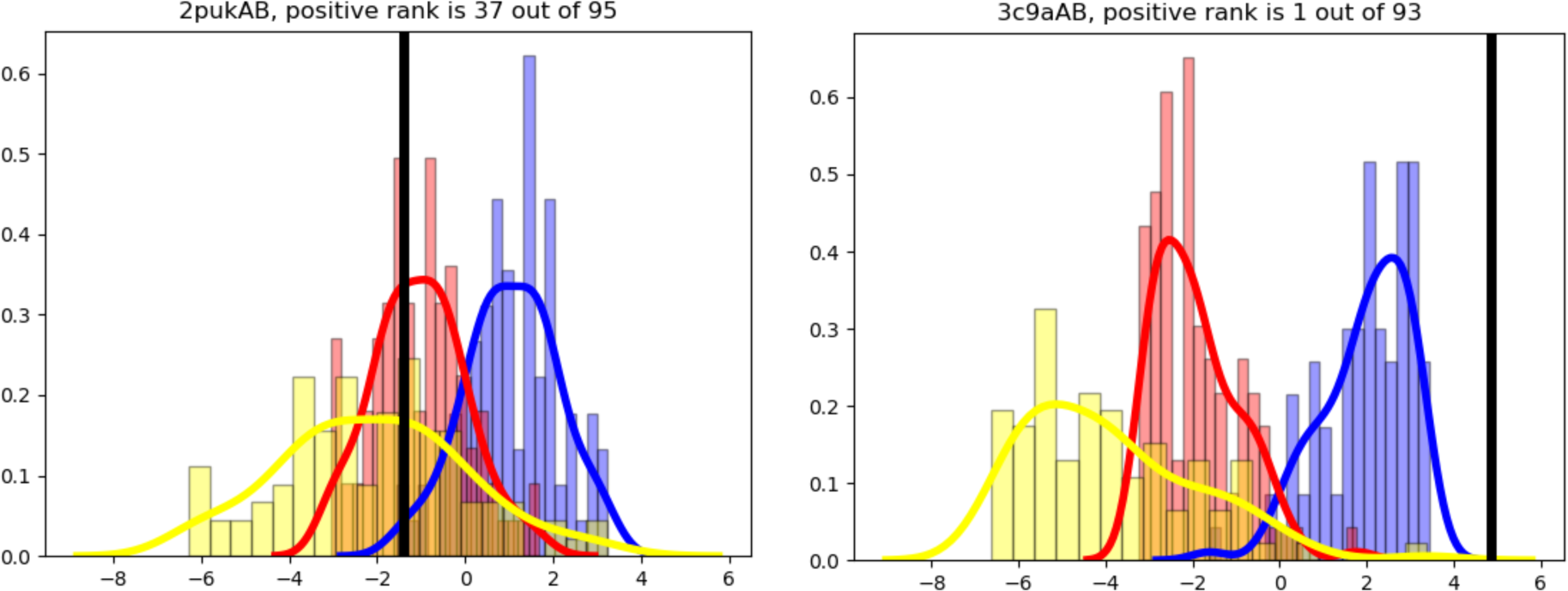
Sample distributions of the CNN scores for two complexes (yellow: diff-score, red: positive, blue: negative).

We further evaluated the performance of our method by determining the rank of the correct interface of a protein-protein complex in the sorted list of all decoys of the same complex. We had only 599 cases from the negative dataset (DOCKGROUND and PPI4DOCK) that has a correct interface in our positive dataset (PIFACE and PDB2012). We ranked all docking poses for each 599 cases and determined the rank of the correct interface. **Table 3** presents the corresponding success rates for the top 1%, 10%, and 20% (i.e, the correct interface being in the top percentage rank). We repeated the study for “training+validation” (454 cases) and “validation+testing” (32 cases) groups. The “validation+testing” group has unseen data. 223 out of 454 (49%) cases have their positives in top 1 in the “training+validation set”. The ratio for the unseen case, drops to 22%, however, still it is considerably high. To put this performance in context, we considered the top percentage rank results from a recent study (CIPS) that ranks protein interfaces (Nadalin and Carbone, 2018). It scores interfaces coming from docking decoys and provides the rank of acceptable interfaces. They reported rank percentages for top1% and top20% of DOCKGROUND dataset as 63% and 94%. Our top 1 and top 20 results are 49% and 86% for training, 22% and 59% for testing. We have to emphasize that the CIPS has 10% near-natives and if any of them is in the top %rank, the complex is flagged as a positive prediction. On the other hand we removed all near native and acceptable models in the decoys and we have only one correct structure (from PDB) which corresponds to 1% of the decoys. With this in mind, the performance of our model can be considered competitive.

## 4. Conclusion

The structures of protein–protein complexes are crucial to better predict new protein complex structures and to understand mechanisms of binding. Identification of protein-protein interactions and interfaces is also at the heart of structure-based drug design, cellular signaling and metabolic pathways. It will also help to understand the effect of disease causing or benign mutations that lie at the protein interfaces. Therefore, it is important to determine how proteins interact through interfaces. DeepInterface, is the first deep learning approach (to our knowledge) to validate a predicted protein-protein interface. It can also be used to rank the interfaces in decoy sets. It only uses atomic coordinates and the amino acid types at the protein interface regions. Our results suggest that DeepInterface is suitable to discriminate incorrect interface structures from the native interface structures. One difficulty that needs to be addressed is the lack of gold-standard negative interface structures. The success rate of methods similar to ours deeply affected by the training datasets. The deep learning approaches promise for more advanced protein-protein interaction studies as more structural data become available.

## References

Adhikari, B., Hou, J. and Cheng, J. DNCON2: improved protein contact prediction using two-level deep convolutional neural networks. Bioinformatics 2018;34(9):1466–1472.

Andreani, J., Faure, G. and Guerois, R. InterEvScore: a novel coarse-grained interface scoring function using a multi-body statistical potential coupled to evolution. Bioinformatics 2013;29(14):1742–1749.

Aytuna, A.S., Gursoy, A. and Keskin, O. Prediction of protein-protein interactions by combining structure and sequence conservation in protein interfaces. Bioinformatics 2005;21(12):2850– 2855.

Baspinar, A., et al. PRISM: a web server and repository for prediction of protein-protein interactions and modeling their 3D complexes. Nucleic Acids Res 2014;42(Web Server issue):W285-289.

Bezanson, J., et al. Julia: A Fresh Approach to Numerical Computing. 2017;59(1):65-98.

Chen, H. and Zhou, H.X. Prediction of interface residues in protein-protein complexes by a consensus neural network method: test against NMR data. Proteins 2005;61(1):21–35.

Chen, R., Li, L. and Weng, Z. ZDOCK: an initial-stage protein-docking algorithm. Proteins 2003;52(1):80–87.

Cukuroglu, E., et al. Non-redundant unique interface structures as templates for modeling protein interactions. PLoS One 2014;9(1):e86738.

He, K., et al. Delving Deep into Rectifiers: Surpassing Human-Level Performance on ImageNet Classification. In, Proceedings of the 2015 IEEE International Conference on Computer Vision (ICCV). IEEE Computer Society; 2015. p. 1026–1034.

He, K., et al. Deep Residual Learning for Image Recognition. In, 2016 IEEE Conference on Computer Vision and Pattern Recognition (CVPR). 2016. p. 770–778.

Ioffe, S. and Szegedy, C. Batch normalization: accelerating deep network training by reducing internal covariate shift. In, Proceedings of the 32nd International Conference on International Conference on Machine Learning - Volume 37. Lille, France: JMLR.org; 2015. p. 448–456.

Janin, J., et al. CAPRI: a Critical Assessment of PRedicted Interactions. Proteins 2003;52(1):2–9.

Jimenez, J., et al. DeepSite: protein-binding site predictor using 3D-convolutional neural networks. Bioinformatics 2017;33(19):3036–3042.

Jones, S. and Thornton, J.M. Analysis of protein-protein interaction sites using surface patches. J Mol Biol 1997;272(1):121–132.

Jordan, R.A., et al. Predicting protein-protein interface residues using local surface structural similarity. BMC Bioinformatics 2012;13:41.

Keskin, O., et al. Empirical solvent-mediated potentials hold for both intra-molecular and intermolecular inter-residue interactions. Protein Sci 1998;7(12):2578–2586.

Keskin, O., et al. Principles of protein-protein interactions: what are the preferred ways for proteins to interact? Chem Rev 2008;108(4):1225–1244.

Keskin, O., Tuncbag, N. and Gursoy, A. Characterization and prediction of protein interfaces to infer protein-protein interaction networks. Curr Pharm Biotechnol 2008;9(2):67–76.

Kingma, D.P. and Ba, J. Adam: A Method for Stochastic Optimization. CoRR 2014;abs/1412.6980.

Kundrotas, P.J., et al. Templates are available to model nearly all complexes of structurally characterized proteins. Proc Natl Acad Sci U S A 2012;109(24):9438–9441.

Lecun, Y., et al. Gradient-based learning applied to document recognition. Proceedings of the IEEE 1998;86(11):2278–2324.

Lensink, M.F., et al. Prediction of homoprotein and heteroprotein complexes by protein docking and template-based modeling: A CASP-CAPRI experiment. Proteins 2016;84 Suppl 1:323–348.

Li, X., et al. Protein-protein interactions: hot spots and structurally conserved residues often locate in complemented pockets that pre-organized in the unbound states: implications for docking. J Mol Biol 2004;344(3):781–795.

Liang, S., et al. Consensus scoring for enriching near-native structures from protein-protein docking decoys. Proteins 2009;75(2):397–403.

Liu, S., Gao, Y. and Vakser, I.A. DOCKGROUND protein-protein docking decoy set. Bioinformatics 2008;24(22):2634–2635.

Liu, Y., et al. Enhancing Evolutionary Couplings with Deep Convolutional Neural Networks. Cell Syst 2018;6(1):65–74 e63.

Liu, Y., et al. Learning structural motif representations for efficient protein structure search. Bioinformatics 2018;34(17):i773–i780.

Nadalin, F. and Carbone, A. Protein-protein interaction specificity is captured by contact preferences and interface composition. Bioinformatics 2018;34(3):459–468.

Nair, V. and Hinton, G.E. Rectified linear units improve restricted boltzmann machines. In, Proceedings of the 27th International Conference on International Conference on Machine Learning. Haifa, Israel: Omnipress; 2010. p. 807–814.

Ogmen, U., et al. PRISM: protein interactions by structural matching. Nucleic Acids Res 2005;33(Web Server issue):W331–336.

Popov, P. and Grudinin, S. Knowledge of Native Protein-Protein Interfaces Is Sufficient To Construct Predictive Models for the Selection of Binding Candidates. J Chem Inf Model 2015;55(10):2242–2255.

Torng, W. and Altman, R.B. 3D deep convolutional neural networks for amino acid environment similarity analysis. BMC Bioinformatics 2017;18(1):302.

Tovchigrechko, A. and Vakser, I.A. GRAMM-X public web server for protein-protein docking. Nucleic Acids Res 2006;34(Web Server issue):W310-314.

Tuncbag, N., et al. Architectures and functional coverage of protein-protein interfaces. J Mol Biol 2008;381(3):785–802.

Tuncbag, N., et al. Predicting protein-protein interactions on a proteome scale by matching evolutionary and structural similarities at interfaces using PRISM. Nat Protoc 2011;6(9):1341–1354.

Winter, C., et al. Protein interactions in 3D: from interface evolution to drug discovery. J Struct Biol 2012;179(3):347–358.

Xavier, G. and Yoshua, B. Understanding the difficulty of training deep feedforward neural networks. In.: PMLR; 2010. p. 249–256.

Xu, Q. and Dunbrack, R.L., Jr. The protein common interface database (ProtCID)--a comprehensive database of interactions of homologous proteins in multiple crystal forms. Nucleic Acids Res 2011;39(Database issue):D761–770.

Yu, J. and Guerois, R. PPI4DOCK: large scale assessment of the use of homology models in free docking over more than 1000 realistic targets. Bioinformatics 2016;32(24):3760–3767.

Yuret, D. Knet: beginning deep learning with 100 lines of Julia. In, Machine Learning Systems Workshop at NIPS 2016. 2016.

Zhao, N., et al. Feature-based classification of native and non-native protein-protein interactions: Comparing supervised and semi-supervised learning approaches. Proteomics 2011;11(22):4321–4330.

